# An oblique plane microscope for mesoscopic imaging of freely moving organisms with cellular resolution

**DOI:** 10.1101/2022.07.15.500249

**Authors:** Rajwinder Singh, Kaushikaram Subramanian, Rory M. Power, Alexandre Paix, Aissam Ikmi, Robert Prevedel

## Abstract

Several important questions in biology require non-invasive and three-dimensional imaging techniques with appropriate spatiotemporal resolution that permit live organisms to move in an unconstrained fashion over an extended field-of-view. While selective-plane illumination microscopy (SPIM) has emerged as a powerful method to observe live biological specimens at high spatio-temporal resolution, typical implementations often necessitate constraining sample mounting or lack the required volumetric speed. Here, we report on an open-top, dual-objective oblique plane microscope (OPM) capable of observing millimeter sized, freely moving animals at cellular resolution. We demonstrate the capabilities of our mesoscopic OPM (MesOPM) by imaging the behavioural dynamics of the sea anemone *Nematostella vectensis* over 1.56 × 1.56 × 0.25 mm at 1.5 × 2.8 × 5.3µm resolution and 0.5Hz volume rate.

## 1. Introduction

In animals, organismal behaviors such as motility and body contraction are integral to their life-cycles, and it is well-established that morphology determines the repertoire of such behaviors. However, it remains unclear how such a link between animal behavior and morphology is established during development [1,2]. This lack of knowledge is in part due to technical challenges of studying organismal biology at a resolution that would enable uncovering mechanistic links between organism size, shape and behavior to be uncovered. In order to bridge the multiple length and time scales of these processes, the sea anemone *Nematostella vectensis* has emerged as an important new model organism in evolutionary developmental biology. In this organism, active muscular hydraulics that control body movement also drive the development of the polyp form [3]. However, several important questions, e.g. how the animal’s behavior, which is driven by muscle hydraulics, guides the animal’s morphology, or how neuronal dynamics control muscle movements and body deformations that are necessary for animal development, remain unanswered.

Because of its high sensitivity to light, relatively large size (∼mm), and constant need to move freely to properly develop and survive, *in toto* 3D live imaging of *Nematostella* with cellular resolution has remained impossible. Although advanced 3D microscopy methods such as selective plane illumination microscopy (SPIM) [4] or light-field microscopy (LFM) [5–8] exist, they typically fail to provide either the necessary spatio-temporal resolution, effective field-of-view, or require the samples to be physically immobilized in order to keep them within the coverage of the microscope’s objective. This is especially true for light-sheet microscopes, as they traditionally rely on separate orthogonal illumination and detection lenses, which complicate sample mounting, require long WD lenses for large FOV, and potentially suffer from light attenuation when going through large tissue samples [9,10]. More recently, so-called single-objective light-sheet microscopes, also known as oblique plane microscopes (OPM) [11,12], have circumvented these geometrical constraints by realizing light-sheet illumination and fluorescence detection from the same objective lens, reducing steric issues for sample mounting all while retaining the advantages of the light-sheet approach in terms of speed, FOV, resolution and gentleness [13]. OPM employs a remote focusing scheme [14] comprising three microscopes in series, for aberration-free re-imaging of an oblique plane of the object into the remote space between the second and third microscopes. A tilted tertiary objective lens then provides a suitably magnified view of the relayed object. By illuminating the oblique focal plane with a suitable light sheet, high-contrast volumetric images can be captured either by illumination/detection scan/de-scan or by stage-scanning the sample.

Despite a large and growing number of recent OPM developments [15,16] also including so-called open-top systems [17–20], it has been challenging to combine these approaches with large FOV, and thus typically low NA, objectives. This is because the necessary re-imaging of the oblique intermediate image plane in OPMs entails substantial optical losses as a large fraction of the light cone projected by the secondary objective lens will lie outside of the acceptance angle of the tilted tertiary objective lens used for imaging [21]. In particular, for NA<0.5 no light will be collected by the third objective, which has so far limited to FOVs to OPMs of ≤1×1mm^2^ [15–20,22]. A notable exception is Ref. [21] which employs a diffraction grating to allow oblique reimaging for low-NA objectives, thereby achieving a large FOV of 3.3 × 3.0mm^2^. However, since this technique utilized a low magnification objective with NA<0.3, the light sheet launch angle is small (17.45 degrees). Since the system PSF is a product of illumination and detection contributions, this small crossing angle yields an axially elongated PSF and poor axial resolution of >30µm - which is not sufficient to discriminate individual cells of most model organisms which tend to be in the range of 5-15µm. Another recent approach sought to artificially increase the light sheet launch angle by employing a grating in the sample space. However, since emitted fluorescence must also traverse the grating, light-losses and aberrations are apparent [23].

Here, we report the design of an OPM specifically tailored to the needs of extended, ‘mesoscopic’ FOVs over 1mm in size, while retaining cellular resolution, i.e. axially ∼5µm. Together with an open-top geometry it enables easy sample handling and the compatibility with imaging of live, freely moving *Nematostella* specimens. We achieve this by adopting a so-called non-orthogonal, dual-objective (NODO) OPM approach [18] which utilizes a separate illumination objective as for traditional SPIM but oriented such that the crossing angle with the detection objective axis is much smaller than 90 °. Since in this configuration the illumination objective is located entirely below the sample plane, the steric hindrance of typical orthogonal light sheet systems is eliminated. We extend the NODO concept to allow for all-optical volumetric imaging, free from requirements of stage scanning for extended FOVs, optimize the illumination angle to minimize losses during the re-imaging step and attenuation of the illumination light through tissue, while maximizing the field of view through careful optical design and a custom tube lens using only commercial off-the-shelf components. We then demonstrate the performance of our microscope by capturing the morpho-behavioural dynamics of freely moving *Nematostella* in 3D and with ∼5µm axial resolution and ∼1Hz volume rate, sufficient to visualize with cellular resolution the complex tissue dynamics during hydraulic contractions in real-time.

## 2. Methods

### OPM principle

In oblique plane microscopes, the illumination and detection planes are tilted relative to the nominal focal plane of the objective. This therefore requires a special optical arrangement taking the oblique detection plane into account when imaging onto an array detector (camera). For volumetric imaging, the oblique excitation light-sheet is then scanned through the volume while the emitted light is de-scanned onto an oblique intermediate image plane. This re-imaging requires a system which satisfies Sine and Herschel conditions simultaneously to minimize (spherical) aberrations associated with imaging of oblique planes. The oblique intermediate plane is then imaged by a tertiary objective and tube lens onto a camera whose optical axis lies perpendicular to this intermediate plane. It is here where a majority of the light loss occurs in single-objective OPMs.

### Microscope design

The schematic design of our MesOPM is illustrated in Fig. 1. CAD drawings and photos of the microscope are shown in Fig. S1. The microscope offers an ‘open-top’ geometry (Fig 1(a)), which facilitates sample mounting without the steric hindrance of traditional light sheet microscopy. In our case, the sample mount is custom designed (Fig. 1(b)) and employs a fluorinated ethylene propylene (FEP) foil to keep the specimen submerged in artificial seawater while separating this medium from the water immersion chamber that is coupled to the inverted, multi-immersion detection objective (configured for water immersion) (O1, 10X 0.5 NA glyc Nikon MRD71120, field-number 22). The primary objective supports a lateral FOV of 2.2 mm diameter while maintaining a high NA of 0.5 given the low magnification. Note that in this configuration, the sample (*Nematostella*) is free to move around unconstrained in a liquid droplet. The optical layout of the microscope is shown in Fig 1(c). For excitation, a 488nm CW laser is coupled into a single-mode fiber which is then expanded using a telescope before passing through an electrically tunable lens (ETL, Optotune EL-10-30-C). A static light-sheet with a waist of ∼4.5µm and length of ∼472 µm (Fig. S2) is generated using a cylindrical lens (CL, f=40mm, Thorlabs LJ1402L1-A). The light-sheet is then relayed onto the sample using a tube lens (TL, Plössl type-2× Thorlabs AC254-125-A) and an objective lens (I1, 5X 0.14 NA air Mitutoyo M Plan Apo). A galvanometric mirror (GM1, Hans scanner ±22.5°) scans the light-sheet over the sample. The obliquely illuminated plane, lies at an angle of 25° relative to the image plane of the primary objective O1. The back-focal plane (BFP) of O1 is conjugated to the galvo mirror GM2 using two identical lenses TL1 and SL1 (f=200mm, Thorlabs TTL200MP). The galvo mirror is then further conjugated to the BFP of the secondary objective (O2, 10X 0.45 NA air Nikon MRD00105) using SL2 (f=200 mm, Thorlabs TTL200MP) and a custom-designed tube lens TL2 with an effective focal length of 150mm (Fig. S3). The tube lenses TL1 and TL2 are chosen such that their focal lengths match the ratio of the refractive indices at the primary and the secondary objective to satisfy the Sine and Herschel conditions. A tertiary objective (O3, 10X 0.45 NA air Nikon MRD00105) is placed relative to the secondary objective at an angle of 25° which is the same as the light-sheet launch angle with respect to O1 (see Fig 1 (c)). The intermediate oblique image plane is then finally reimaged onto a sCMOS camera (Kinetix, Teledyne Photometrics) using a tube lens (TL3, f=200mm, Thorlabs TTL200A). Since the effective overall magnification of the system is 13.3 and the chip size of the camera 20.8×20.8mm^2^, the overall FOV is ∼1.56×1.56mm (diagonal 2.2mm).

**Fig. 1:**
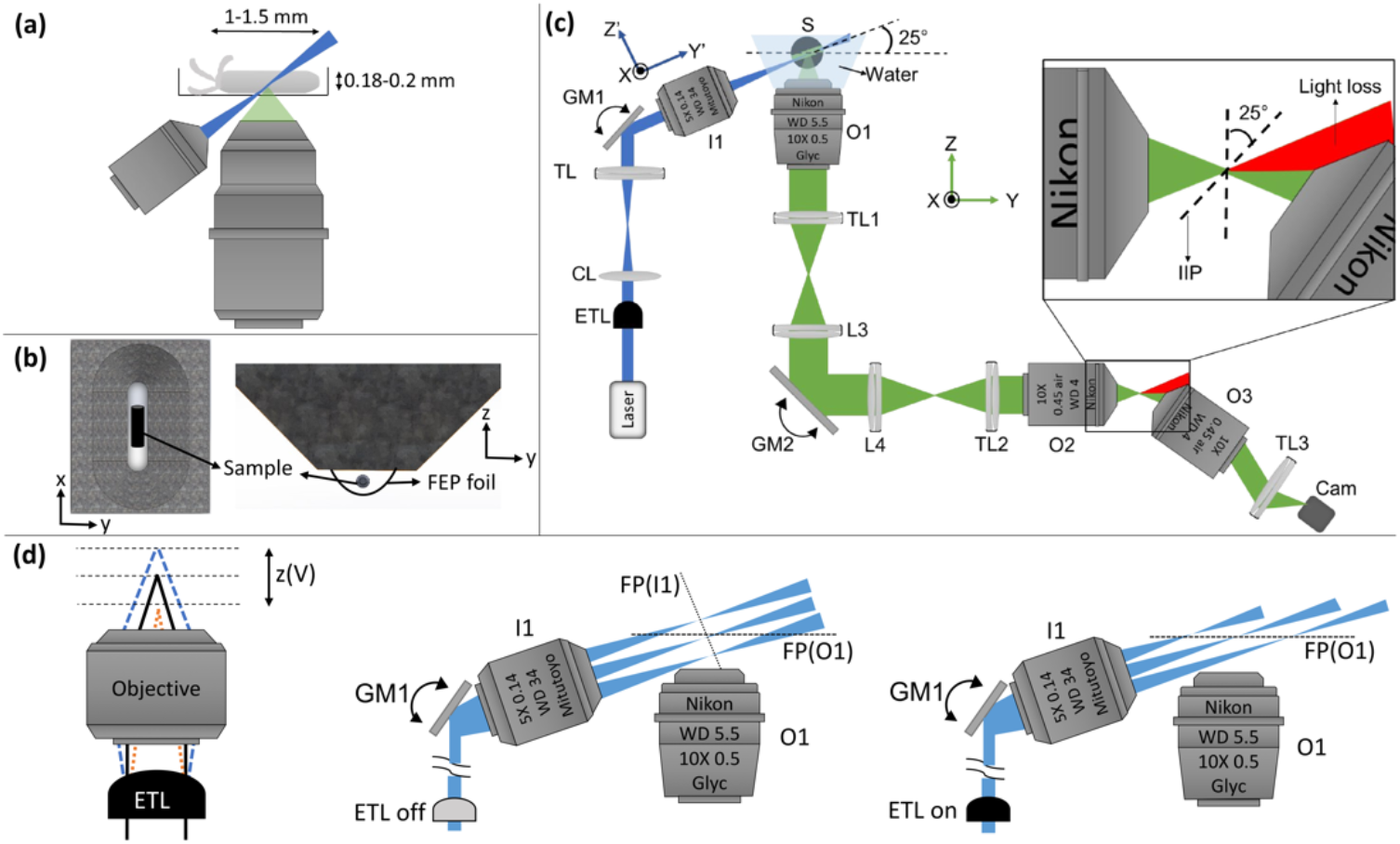
(a) MesOPM geometry (NODO configuration) showing illumination and detection objectives with respect to the sample (Nematostella polyp). (b) Custom sample mount which holds the sample in the imaging chamber separated by a FEP foil. (c) Optical layout of the microscope. The setup has three main units. The illumination objective (I1) launches the light-sheet into the water chamber (WC) through a glass window that lies perpendicular to the propagation axis of the lightsheet (also see Fig. S1). The detected signal is then collected by the detection objective (O1) and relayed at the focus of the secondary objective (O2), IIP -intermediate image plane. The image is de-magnified by 1.33 times to satisfy the Sine and Herschel condition. The focal plane of the tertiary objective (O3) coincides with the tilted image which is then imaged onto the camera. The proportion of the light being lost due to the tilt of O3 with respect to O2 is shown in red. (d) Left: Concept of the electro-tunable lens (ETL). In contrast to scanning a static light-sheet (middle), dynamically refocusing the light-sheet with an ETL (right) ensures the overlap with the focal plane of O1 during volumetric scanning over large lateral FOVs.

### Design rationale

For volumetric imaging, the light-sheet is scanned through the volume via GM1 which is synchronized to GM2 in order to de-scan the fluorescence onto the stationary intermediate image plane (IIP) shown in the zoom-in in Fig. 1(c). This eliminates the need for stage scanning, allowing us to volumetrically image the sample using a galvanometric mirror, which is faster and non-perturbative to the sample by virtue of its remote location and comparably minimal inertia. The microscope uses a unique scan geometry [24] to achieve tilt-invariant light-sheet scanning at the focal plane of the primary objective. An electrically tunable lens (ETL) is synchronized to the galvo scanner (GM1) and is utilized to dynamically adjust the focus of the light-sheet such that its focus coincides with the focal plane of O1 across the scan range, thereby ensuring optimal and constant axial resolution and sectioning throughout the lateral FOV of >1.5mm (Fig 1(d)). The launch angle of 25deg. was chosen to optimize the axial resolution and FOV, while keeping the light-losses between O2-O3 tolerable. As shown in Fig.S4, 25deg. yields an axial FOV of ∼240µm, sufficient to capture an entire *Nematostella* polyp, while only losing ∼46% of the light between O2 and O3.

In order to satisfy the Sine and Herschel conditions in the remote focusing unit, the tube lens of the secondary objective should ideally have a focal length of 150mm. However, suitable widefield tube lenses are not commercially available. We therefore designed a custom tube lens for that purpose that is corrected to attain a field size of 1.5 mm at the object over the relevant pupil size when combined with our 10x objectives (Fig. S3). Furthermore, in order to minimize vignetting effects of the tube lenses in the system when used in telecentric 4f-configurations and retain diffraction-limited performance over the large FOV, we simulated the performance of several commercial tube lenses and found only a single one (Thorlabs, TTL200-MP) to have an aperture large enough to not cause vignetting.

### 3D volume reconstruction

In oblique plane microscopy, a 3D volume is acquired plane-by-plane in a sheared geometry. This must be converted into the correct rectilinear lab coordinate system. To achieve this, an affine transformation has to be applied to the 3D dataset [25]. The illumination plane in our case is tilted relative to the optical axis of O1 by 65 degrees and thereby the affine transformation matrix used to deskew the 3D volumes of our MesOPM becomes:

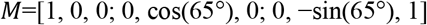

We used the TransformJ plugin available in the ImageJ software to reconstruct the 3D volume acquired on the system, and utilized a voxel size of 0.488 × 0.488 × 1.3μm^3^ of the raw image stack as given by the total magnification of the system (13.3x), pixel size of the camera (6.5µm) and minimum step size of the digital galvo scanner employed (d = 1.3µm).

## 3. Results

### Optical characterization

The optical performance of the microscope was characterized by imaging 0.5 µm diameter TetraSpeck Microbeads suspended in a transparent gelrite solution with light sheet illumination at 488 nm. The resolution was determined quantitatively by calculating the full-width at half maximum (FWHM) of the beads across the entire FOV in the axial and lateral direction after transformation of the 3D stack into the lab coordinate system. Fig. 2 summarizes these findings. Fig 2(a) shows the field corrected z-projection of a 3D stack composed of 200 frames (each 1.3 µm apart in the y-direction) to compensate for the decreasing intensity towards the edges due to the Gaussian profile of the beam. A representative PSFs of the system is shown in Fig. 2(b). The FWHM in the x, y, and z-directions was determined by averaging over 50 beads randomly selected across the FOV and found to be 1.62±0.38µm, 2.81±0.48µm, and 5.27±0.61µm, respectively. Furthermore, the spatial dependence of the resolution across the FOV was calculated by dividing the lateral and axial FOV into 10 equally spaced sections (∼150µm wide) and the FWHM was calculated by averaging 10 randomly selecting beads from each of them (Fig 2(c,d)). The spatial resolution was not found to vary substantially due to the dynamic ETL re-focusing during volume scanning. The resolution of the microscope was also compared to theoretical predictions using a recently published pipeline [18]. Given the overall effective pupil function of the microscope, the FWHM along the x-, y-and z-directions were calculated as 0.62 µm, 1.16 µm, and 4.24 µm, respectively (Fig. S5). Here we note that the experimentally obtained values are limited by the detector pixel size and the scan step of the galvo in the x and y direction respectively. Given the total magnification of the system (13.33x) and the pixel size of the camera (6.5 µm) as well as the scan step of the galvo in the y-direction (∼1.3 µm), the upper bound to the lateral resolution according to the Nyquist criteria is 0.98 µm in x and 2.6 µm in the y-direction. The experimentally obtained values, and here especially the axial resolution, are in very good agreement with the theoretical value, further validating our optical design and realization. The overall performance of our OPM in terms of spatial resolution and FOV compares favorably with previous work in the field (Table S1, Fig. S6).

**Fig. 2:**
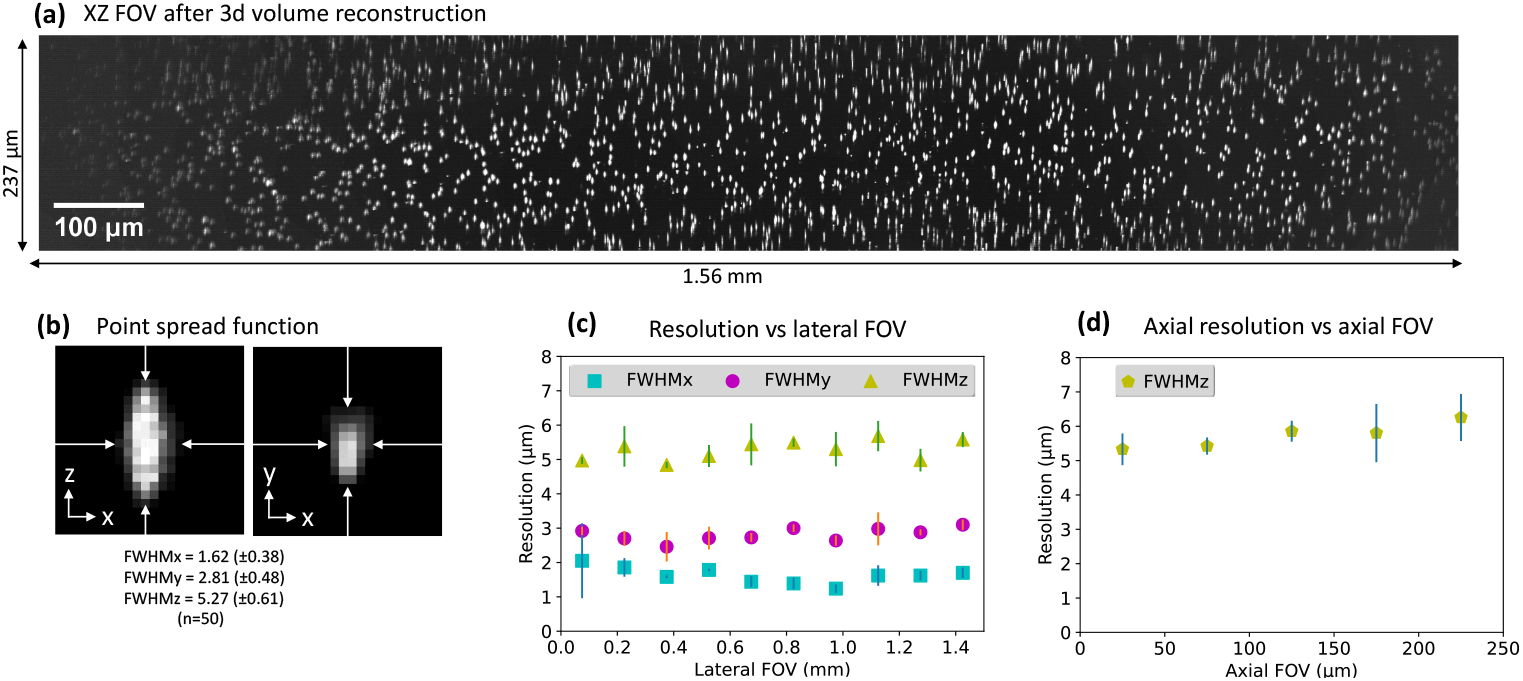
(a) A raw 3D image stack of beads captured over the FOV of 1.56×1.56×0.24mm^3^, projected along the y-axis (xz-plane). (b) Exemplary lateral and axial PSF of a single bead. The FWHM along the x-, y-and z-directions averaged over (n=50) beads are 1.62±0.38µm, 2.81±0.48µm, and 5.27±0.61µm, respectively. The resolution is further plotted as a function of the (c) lateral, and (d) axial FOV.

### Nematostella imaging

After characterizing the optical performance of the MesOPM on static beads, we set out to demonstrate its capabilities on freely moving *Nematostella* primary polyps. In particular, the large FOV and high spatial resolution of our MesOPM system make it ideal for imaging cellular dynamics across whole organisms without the need of moving the sample itself. This is especially important in capturing the subject’s intrinsic behaviors with minimal disturbance of the organism. In combination with the relatively simple bi-layered epithelial anatomy of the primary polyp, MesOPM enables the whole-animal scale imaging of its neuromuscular system and behaviors.

We first imaged the muscular structure of one week old anesthetized Nematostella polyps expressing a fluorescently tagged Myosin Heavy Chain protein (MHC-mNeongreen) over the FOV of 1.5mm in large water droplets mounted in our custom sample holder. A maximum projection of an entire FOV composed of 429 individual frames can be seen in Fig. 3 (a) and Media S1. Zooming into the dataset (Fig. 3(b-d)), closer analysis of individual slices reveals both parietal (high intensity) and ring (low intensity) muscles of the body wall (Fig. 3(d)). These two types of muscles develop in the inner body layer and drive body contractions.

**Fig. 3:**
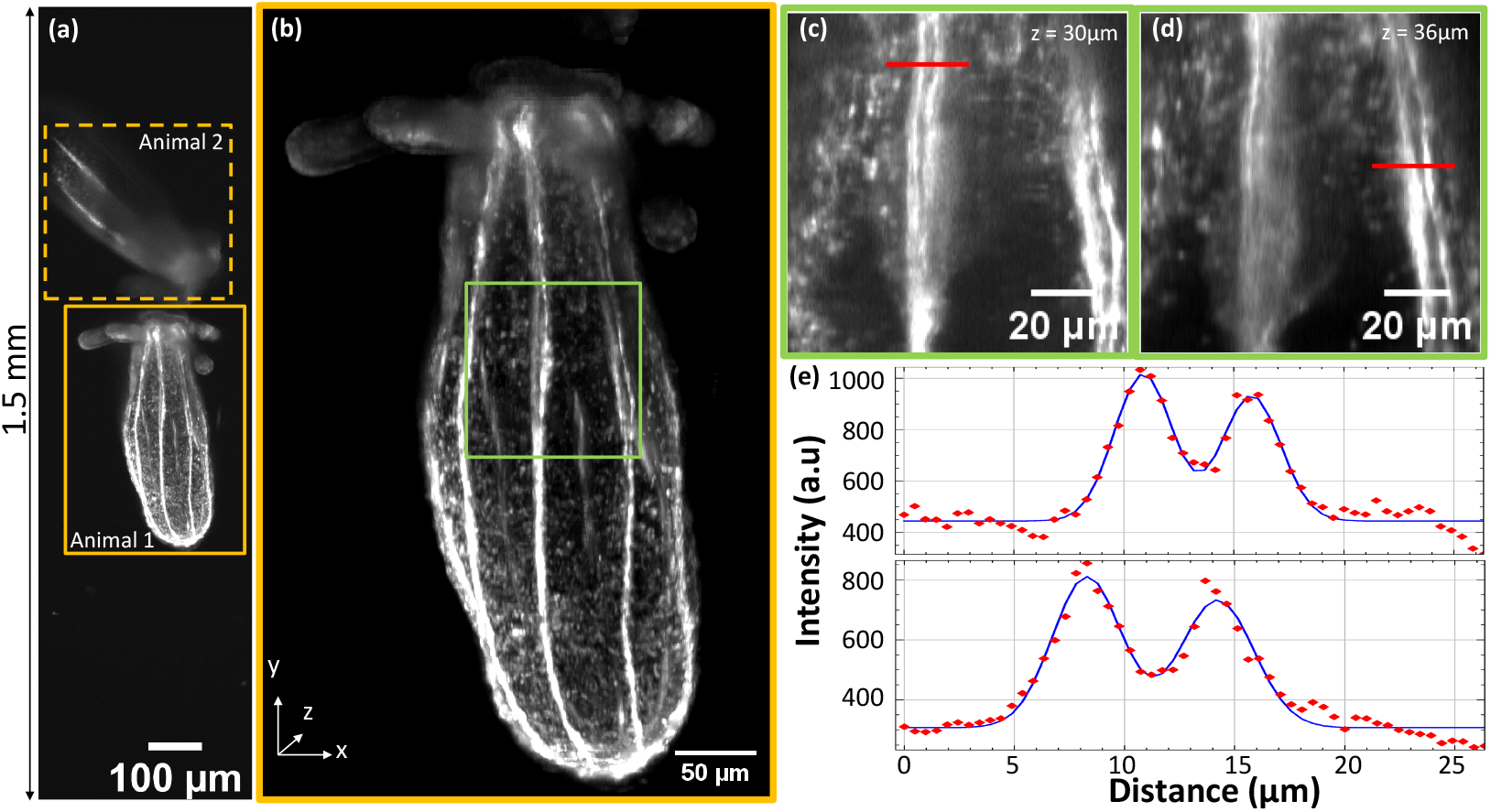
(a) Maximum intensity projection of the imaging volume comprising two *Nematostella* polyp (Myosin Heavy Chain-mNeongreen) imaged on our MesOPM, illustrating the large FOV. The entire stack comprises 429 frames, 1.3 µm apart, 40ms exposure time, 0.5mW excitation power. Total acquisition time for volume: 21 sec. (b) Zoom-in to an individual animal as indicated by the orange box in (a). Also see Media S1. (c-d) Zoom-in of green square in (b) showing parietal muscular structures at two different z-planes. (e) Line intensity plot across muscle fibers.

Next, we imaged freely moving primary polyps expressing Cadherin 1 tagged with eGFP (Cdh1-eGFP) (Fig. 4). We took advantage of the localization of this protein that labels apical adherens junctions to visualize cell-cell contacts in the body wall [26]. A maximum intensity projection (Fig. 4(top)) and zoom-in of XY and XZ planes (Fig. 4(bottom)) clearly show that our MesOPM can identify the cell junctions and thus cellular boundaries. This confirms the high spatial resolution of the microscope and the ability to image with cellular resolution under realistic imaging conditions (laser power 2 mW, exposure time per frame 50 ms) while simultaneously capturing the global organismal tissue level.

**Fig. 4:**
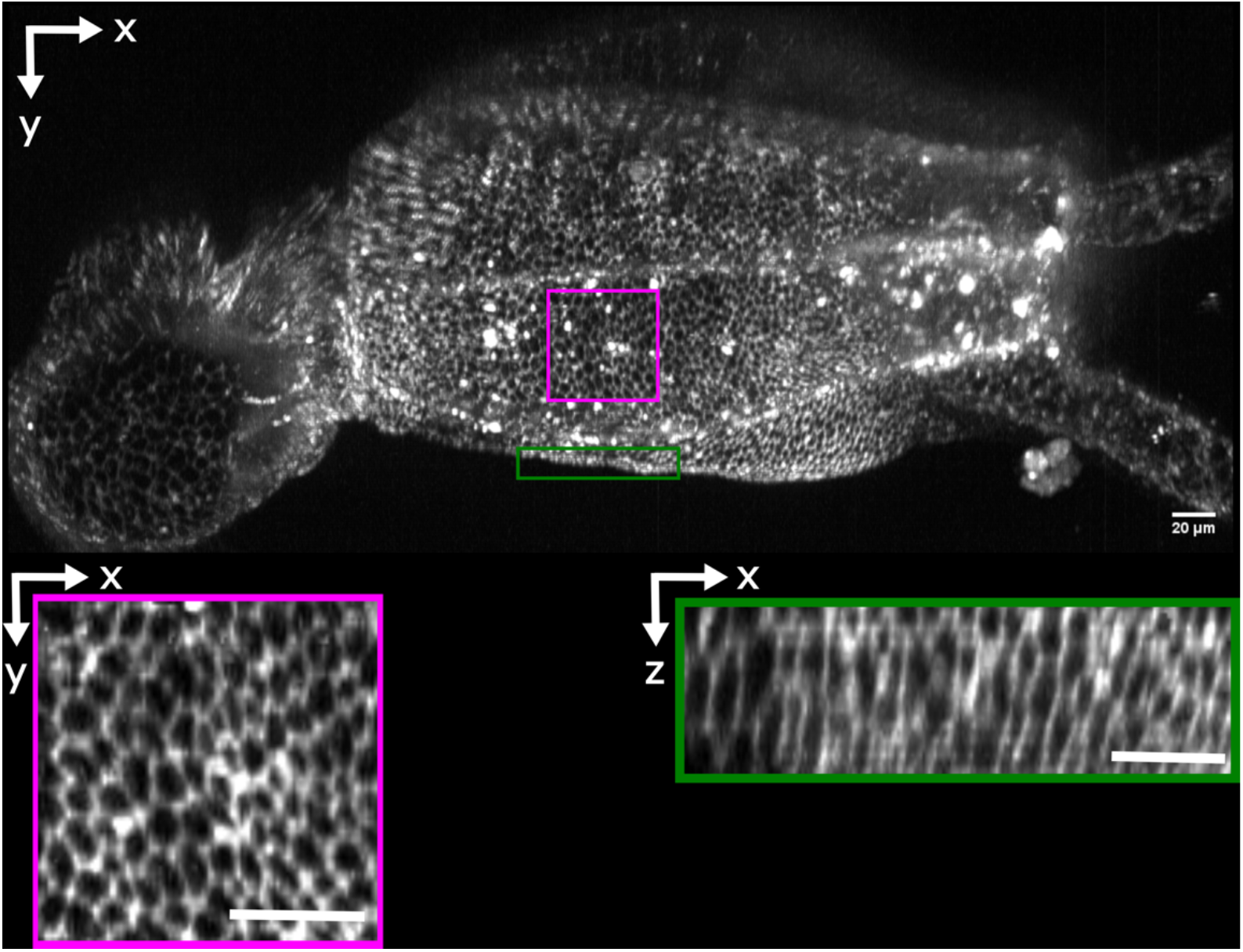
Maximum intensity projection of the imaging volume of *Nematostella* polyp (cadh1-eGFP). The entire stack comprises 500 frames, 1.3 µm apart, 50ms exposure time, 2mW excitation power. Zoom-boxes highlight the high spatial resolution of our MesOPM both in the lateral (magenta box) as well as axial (green box) dimension. Cellular outlines are clearly resolved in both, confirming the high spatial resolution of the microscope in the lateral and axial dimension under realistic imaging conditions.

We then set out to demonstrate the achievable volumetric imaging speed while fully capitalizing on the large optical FOV of our MesOPM. For this we chose a high-speed, large-sensor sCMOS camera (3200×3200 pixel, Kinetix sCMOS, Teledyne Photometrics) in order to be able to capture the nerve net of the polyp during a peristalic wave. Fig. 5 and Media S2 clearly show the distribution of the neurons along the body wall at a resolution that enables observing their cell bodies and fine neurites despite body movement. By acquiring 300 frames per second, we were able to volumetrically image the FOV that contained the entire polyp in ∼2 sec.

**Fig. 5:**
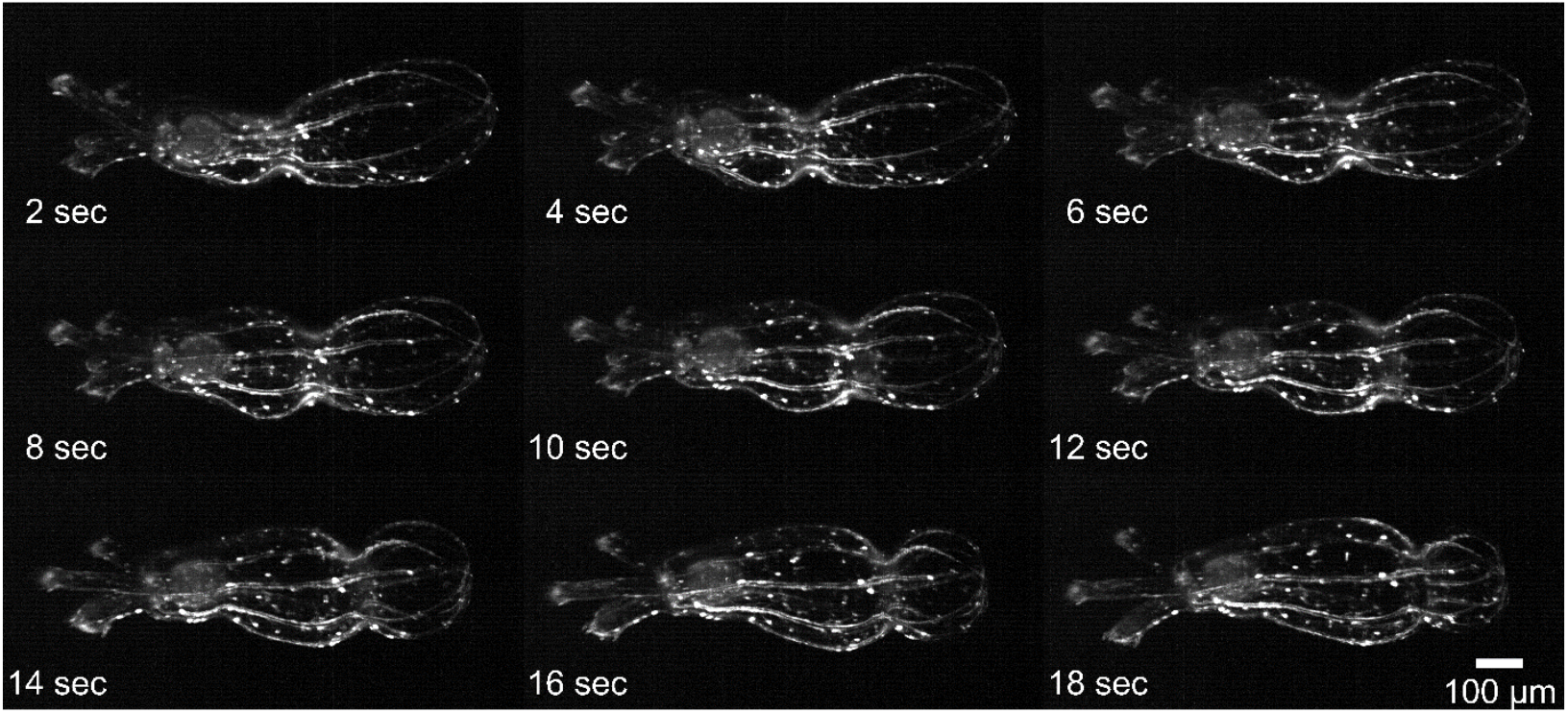
Time series of Nematostella polyp (Elav>mb-eGFP) undergoing body contraction along oral-aboral axis. Each image is a cropped 3D projection of a volume (1.56*0.78*0.24 mm^3^) acquired. The total laser power at the sample was 2mW and the acquisition time 1ms per plane. Each volume is composed of 600 frames acquired at 300fps, yielding a volume every 2 seconds (0.5Hz). Also see Media S2.

## 4. Discussion

To summarize, in this work we reported the first OPM design for mesoscopic scales above 1.5mm FOV that retains cellular resolution (i.e. ∼5µm axially) without the need for stitching multiple volumes together or elaborate sample movement. We used the microscope to perform the first fluorescence 3D live imaging of Nematostella polyps *in-toto* during natural behavior. This establishes our MesOPM as the first imaging apparatus that allows to simultaneously map morpho-behavioral dynamics and neural activity of aquatic invertebrates in 3D.

From a technical point of view, our OPM compares favorably with other, recently published systems, as detailed in Table S1 and Fig. S6. In particular, it provides a substantially larger FOV compared to Ref. [15,16], albeit at the expense of slightly worse spatial resolution. At the moment, only Refs. [21,27] provide much larger accessible image volumes, however we note that these approaches yield a substantially worse resolution (∼30µm and 15µm axially, respectively), which would be insufficient for most cellular imaging experiments. Here the combination of our optical design with a large area and high-frame rate sCMOS camera fully unlocks the potential of the MesOPM for capturing the dynamics of freely moving animals. Finally, we note that another recent mesoscopic OPM approach that has comparable overall imaging performance was recently put forward, which utilizes a transmission grating in the object space to generate the oblique light-sheet [23]. In contrast to our approach the space occupied by the grating limits the available working distance, precluding combination with higher NA objective lenses, while the fluorescence emission needs to also traverse the grating, introducing additional losses along with chromatic and spherical aberrations.

The main limitation of our current system is shared with other light-sheet implementations, namely the degradation of the signal brightness and contrast at the sample top, i.e., facing away from the objective lenses delivering illumination and collecting the resulting fluorescence. In this regard, absorption as well as refraction and scattering resulting from bulk and small-scale variations in refractive index cause an attenuation or broadening of the light sheet and a loss of positional information of the individual emitters deep into tissue, thus resulting in a depth dependent loss of image quality. In principle, these effects could be mitigated by the use of adaptive illumination schemes, such as fast axial sweeping and power modulation or pre-compensation for depth-dependent attenuation of the light sheet [28], real-time active-alignment control [29] or refractive index matching approaches [30]. Furthermore, the reported imaging speed does not fully leverage the available readout and triggering modes of the camera offering the potential for > 2x increases in the volumetric imaging rate. Multiplexed spatially distributive imaging schemes utilizing multiple cameras offer a potential route to greater imaging speed still [31].

Going forward, our MesOPM will now allow us to experimentally address fundamental questions in developmental biology such as how the interplay between morphogenetic, behavioral and neurobiological processes impact the development of animal form and function. Combined with a growing number of transgenic reporter lines for biological processes, we expect that our MesOPM will become a powerful tool to provide further insights into the functional design principles of animal development and behavior.

## Supporting information

Media S1

Media S2

## Funding

This work was supported by funds from the European Molecular Biology Laboratory. K.S. is supported by a research fellowship from the EMBL Interdisciplinary Postdoc (EIPOD) Program under Marie Skłodowska Curie Cofund Actions MSCA-COFUND-FP (664726).

## Acknowledgements

We would like to acknowledge support by the EMBL Heidelberg mechanical and electronic workshops, especially Alejandro Gil Ortiz, as well as Lars Hufnagel, Anniek Stokkermans and Sebastian Hambura for help and support. We are further grateful for useful advice from Adam Glaser during the initial phase of our project.

## Disclosures

The authors declare that there are no conflicts of interest related to this article.

## Data availability

Data underlying the results presented in this paper are not publicly available at this time but may be obtained from the authors upon reasonable request.

## Notes

### Competing Interest Statement

The authors have declared no competing interest.

