## Supplementary Information for "An oblique plane microscope for mesoscopic imaging of freely moving organisms with cellular resolution"

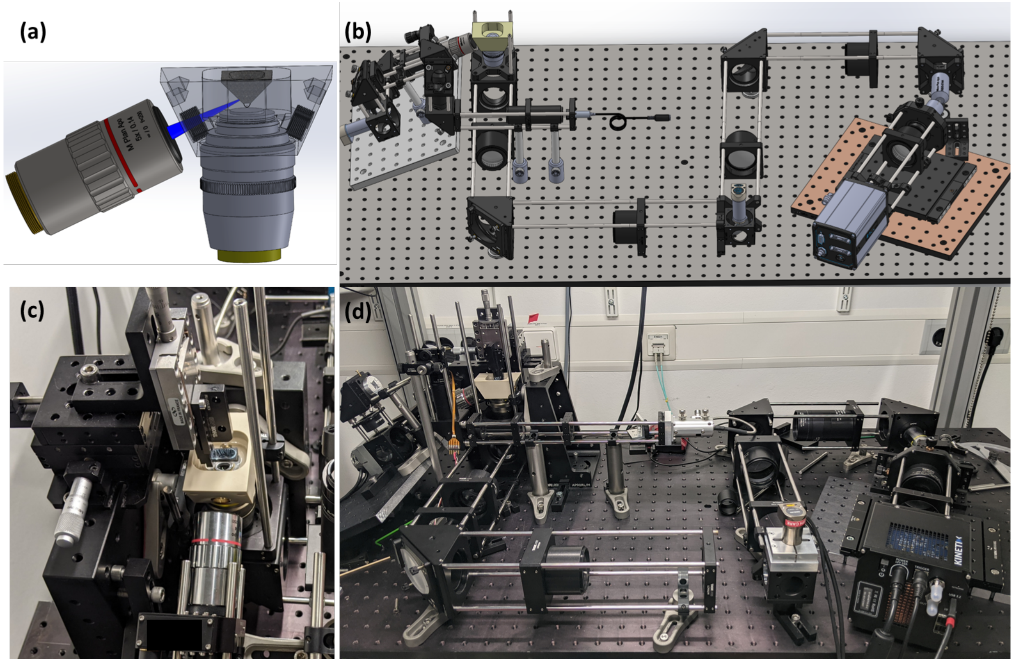


**Figure S1: Setup drawings and implementation.** CAD rendering showing (a) the sample mounting as well as (b) the entire MesOPM setup. The light-sheet is launched into the water chamber through a glass window that sits perpendicular to the propagation axis of the light-sheet. (c) Photo of the sample mounting stage. The sample is loaded from the top and rests on a FEP foil. (d) Photo of the entire MesOPM optical setup.

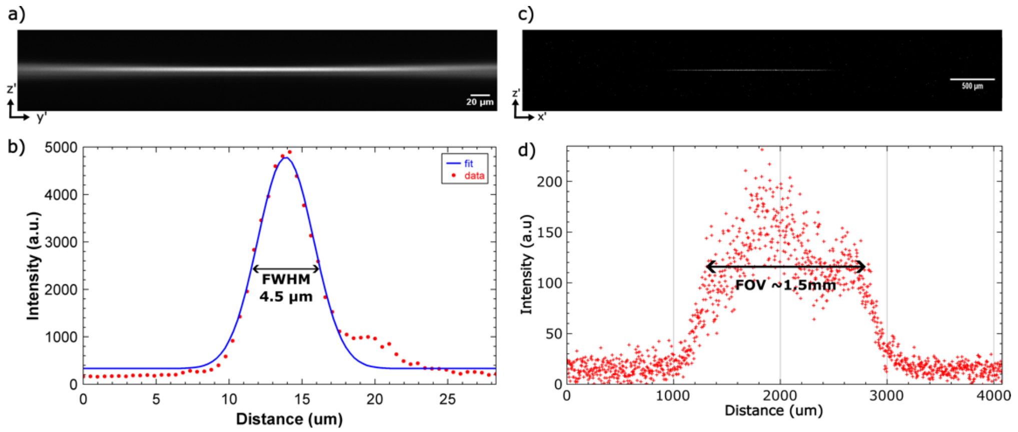


**Figure S2: Excitation light-sheet characterization.** (a) Profile of the Gaussian excitation beam across the (axial) FOV. The beam diameter (FWHM) at the center is plotted in (b). (c-d) Characterization of the light-sheet height, showing grey values plotted over the FOV, indicating sufficient excitation intensity for the 1.5mm FOV of the system.

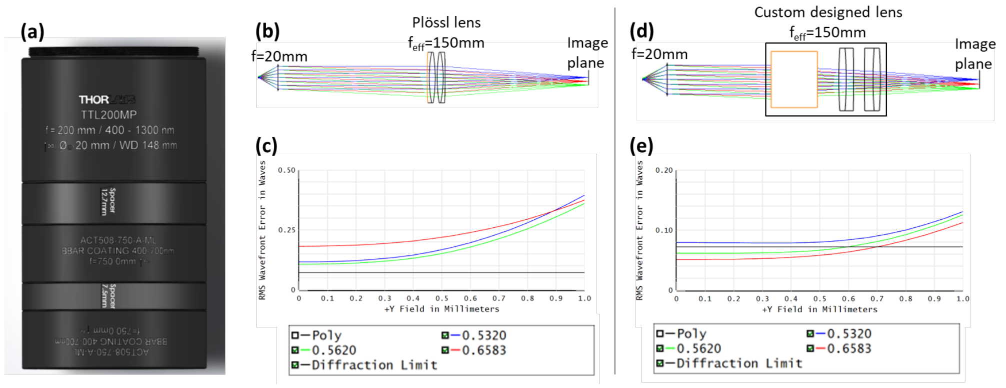


**Figure S3: Custom-designed tube lens.** (a) CAD model of the custom tube lens constructed from off-the-shelf components. (b) A Ploessl lens model with effective focal length of 150mm used as a tube lens in combination with a paraxial lens of focal length 20 mm to image a FOV of 2 mm onto the image plane. (c) RMS wavefront error over the FOV for different wavelengths. The wavefront error remains well above the diffraction limit shown as black straight line. (d) Our custom designed lens used in combination with a paraxial lens of focal length 20mm. (e) RMS wavefront error remains close to the diffraction limit over the whole FOV. Thus, the optical performance of our custom-designed lens is superior to a conventional Ploessl lens configuration.

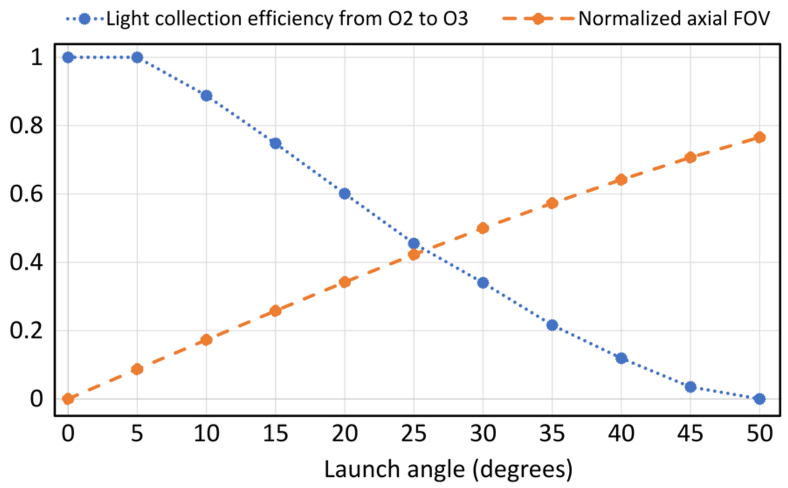


**Figure S4: Effect of light-sheet launch angle on axial FOV and light efficiency.** (a) The theoretical light collection efficiency and the normalized axial FOV of our system are plotted as the function of light-sheet launch angle. In our case 25 degrees was determined as the optimal light-sheet launch angle for our system given the requirements on axial FOV.

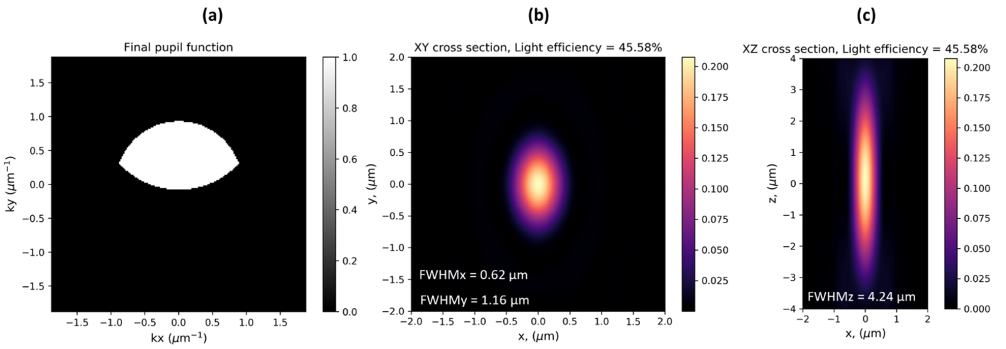


**Figure S5: Theoretical modeling of the PSF of our system.** (a) The effective pupil function of the system. (b) Lateral resolution of the system calculated from the pupil function. The theoretical FWHM in x-direction is 0.62 μm and 1.16 μm in y-direction. (c) The theoretical axial resolution (FWHM in z-direction) of the system is 4.24 μm. Simulation based on Ref. [18].

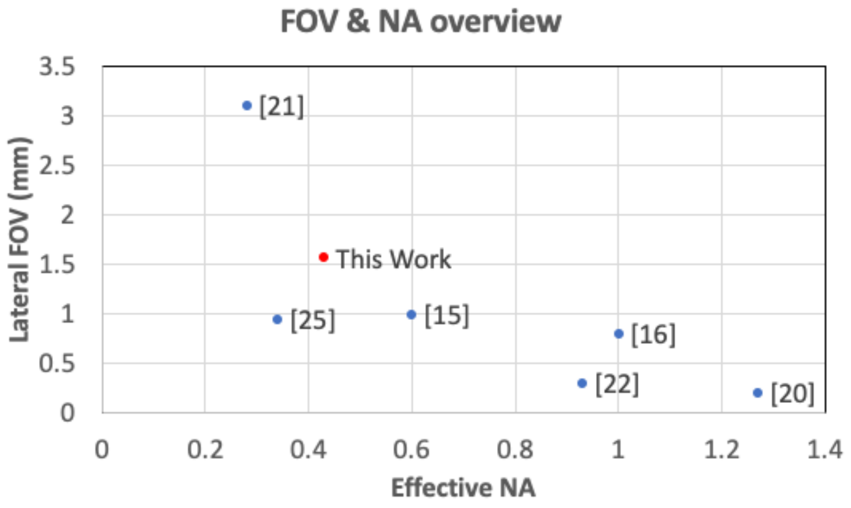


**Figure S6: FOV and effective NA of recent state-of-the-art OPM systems.** References in main manuscript are given in [brackets].

| **Technique [Reference]** | **SCAPE 2.0 [15]** | **DaXi [16]** | **SOPi [25]** | **eSPIM [20]** | **DOPM [21]** | **dOPM [22]** | **Our system** |
|---|---|---|---|---|---|---|---|
| **Effective detection NA** | 0.60*0.35 | 1.0*0.97 | 0.34*0.6 | 1.27*1.18 | 0.28*0.28 | 0.93*0.58 | 0.43*0.23 |
| **Lateral resolution (x*y µm)** | 1.2*0.6 | 0.4*0.4 | 1.3*1.3 | 0.39*0.31 | 3.3*3.0 | 0.39*0.35 | 1.6*2.8 |
| **Axial resolution (µm)** | 2 | 2 | Not specified | 0.6 | 37 | 0.81 | 5.3 |
| **Lateral FOV (x*y mm)** | 1.0*1.0 | 0.8*0.8 | 0.95*0.32 | 0.7*0.7 | 3.1*2.6 | 0.3*0.3 | 1.56*1.56 |
| **Axial FOV (mm)** | 0.38 | 0.3 | 0.4 | 0.2 | 1 | 0.2 | 0.24 |

**Table S1: Performance comparison of recent state-of-the-art published OPM systems.** References in main manuscript are given in [brackets].

**Media S1: 3D rendering of Nematostella polyp (Myosin Heavy Chain-mNeongreen) imaged on our MesOPM.** The entire volume comprises 429 frames, 1.3 µm apart, 40ms exposure time, 0.5mW excitation power. Total acquisition time for volume: 21 sec. Also see Fig. 3.

**Media S2: Video of Nematostella polyp (Elav>mb-eGFP) undergoing body contraction along oral-aboral axis.** Each volume is composed of 600 frames acquired at 300fps, yielding a volume every 2 seconds (0.5Hz). The total laser power at the sample was 2mW and the acquisition time 1ms per plane. Also see Fig. 5.
